# PINAWeb: A web-based tool for the comparison of Protein-Protein Interaction Networks Aligners

**DOI:** 10.1101/2021.04.23.441154

**Authors:** A. Alcalá, G. Riera, I. García, R. Alberich, M. Llabrés

## Abstract

**Motivation:** Several protein-protein interaction networks (PPIN) aligners have been developed during the last 15 years. One of their goals is to help the functional annotation of proteins and the prediction of protein-protein interactions. A correct aligner must preserve the network’s topology as well as the biological coherence. However, this is a trade-off that is hard to achieve. In addition, most aligners require a considerable effort to use in practice and many researchers must choose an aligner without the opportunity to previously compare the performance of different aligners.

**Results:** We developed PINAWeb, a user-friendly web-based tool to obtain and compare the results produced by the aligners: AligNet, HubAlign, L-GRAAL, PINALOG and SPINAL. PPINs can be uploaded either from the STRING database or from a user database. The source code of PINAWeb is freely available on GitHub to enable researchers to add other aligners, network databases or alignment score metrics. In addition, PINAWeb provides a report with the analysis for every alignment in terms of topological and functional information scores, as well as the visualization of the alignments’ comparison (agreement/differences) when more than one aligner are considered.

**Availability:** https://bioinfo.uib.es/~recerca/PINAWeb

## 1 Introduction

Predicting new protein functions and protein-protein interactions is a relevant topic in computational biology (Rost, Liu, Nair, Wrzeszczynski, & Ofran, 2003; Sleator & Walsh, 2010; Kulmanov & Hoehndorf, 2019; Jiang et al., 2016). This topic has played a key role during the COVID-19 pandemic to find out a vaccine for the coronavirus disease (Gordon et al., 2020; Khorsand, Savadi, & Naghibzadeh, 2020; M & G., 2020). Indeed, understanding the mechanism of viral infections is a crucial step towards the discovery of antiviral drugs and vaccines. In this line of research, virus-host protein-protein interaction networks, a particular form of protein-protein interaction networks (PPINs), have become appropriate to analyze virus-host relationships. Specifically, information on well-known and studied virus-host protein-protein interaction networks can be transferred to new ones through protein-protein interaction network comparison and alignment. The same rule applies in the general setting of protein-protein interaction networks, where biological knowledge of well-known and studied protein-protein interaction networks is transferred to new ones through a network alignment. A popular approach in this framework is the “network-based” approach (Sharan, Ulitsky, & Shamir, 2007). With this approach, new interactions between proteins are understood as “missing links” in protein-protein interaction networks. New functional information is obtained either from the characteristics of the protein’s position in the network or it is transferred from annotated proteins sharing a similar position in the PPIN of other organisms. To transfer functional information between proteins from different organisms it is necessary to relate the proteins sharing a similar position in the PPINs. This relation between proteins sharing a similar position is obtained by a pairwise PPINs alignment. Within this alignment setting, aligners with a high ratio of preserved edges match proteins that share a similar position in the networks. On the other hand, aligners with high functional coherence values match proteins sharing some biological function. Hence, the following question arise: which is the best aligner one can use to achieve the prediction of new protein functions and protein-protein interactions via a PPINs alignment?

Many PPINs aligners have been developed along the last decade (Alcalá, Alberich, Llabrés, Rosselló, & Valiente, 2020; Hashemifar & Xu, 2014; Malod-Dognin & Pržulj, 2015; Phan & Sternberg, 2012; Aladağ & Erten, 2013) as well as several scores to assess its performance. The correctness of an alignment is measured by the preservation of the network structure and the proteins functional information. See surveys (Ali, Viswanath, Patil, & Venugopal, 2017; Clark & Kalita, 2014; Elmsallati, Clark, & Kalita, 2016; Ma CY, 2020; Erten, 2021) for a review of implemented aligners and their evaluation. However, it is very difficult to fulfill both requirements. Indeed, the authors in (Clark & Kalita, 2014) conclude that:“*we find dramatic differences between existing algorithms in the quality of the alignments they produce. Additionally, we find that many of these tools are inconvenient to use in practice, and there remains a need for easy-to-use, cross-platform tools for performing network alignment”*. In addition, the authors in (Ma CY, 2020) state that: “*in the network alignment problem, unfortunately there is no gold standard for evaluation; that is, a best alignment is unknown from a biological perspective”*. Therefore, when the topological similarity between two networks wants to be captured rather than the biological function similarity of the matched proteins, then an alignment tool with a high ratio of edges preserved by the alignment must be considered. On the other hand, when the function similarity of the matched proteins is more relevant than the edge preservation, then a suitable alignment tool is an aligner providing a high functional coherence score.

This current lack of a “one size fits all” PPIN aligner together with the computational problems that arise when installing their implementations, motivated us to develop a user-friendly web-based tool to obtain and compare the results produced by different aligners. Thus, we have devised PINAweb, a web based tool to perform a pairwise alignment of two PPINs, considering the aligners: AligNet (Alcalá et al., 2020), HubAlign (Hashemifar & Xu, 2014), L-GRAAL (Malod-Dognin & Pržulj, 2015), PINA-LOG (Phan & Sternberg, 2012) and SPINAL (Aladağ & Erten, 2013), which are the most recent and the best evaluated in (Clark & Kalita, 2014). For every aligner and pair of networks, PINAWeb returns the corresponding alignment, its topological and biological correctness scores and also the visualization of the alignments comparison (agreement/differences) when several aligners are considered. In addition, being that running an aligner on its own data is a major problem for most researchers, PINAWeb has been devised such that PPINs can be uploaded either from a standard database or from the researcher’s own data. Finally, it is an open tool, and new aligners, new databases, and new metrics for scoring alignments can be integrated.

## Implementation

PINAWeb is organized under the philosophy of microservices instead of a monolithic architecture (Newman, 2015). The main benefit of such a design is that each small piece is responsible of only one type of task. In this way, each piece of the tool has no dependencies caused by other pieces and it is easier to detect errors. As shown in Figure 1, PINAWeb’s architecture is divided into four main layers: the front-end, the results dashboard, the REST API, and the server. Then, we use connectors to link these layers. The main information flow is the following:

**Figure 1:**
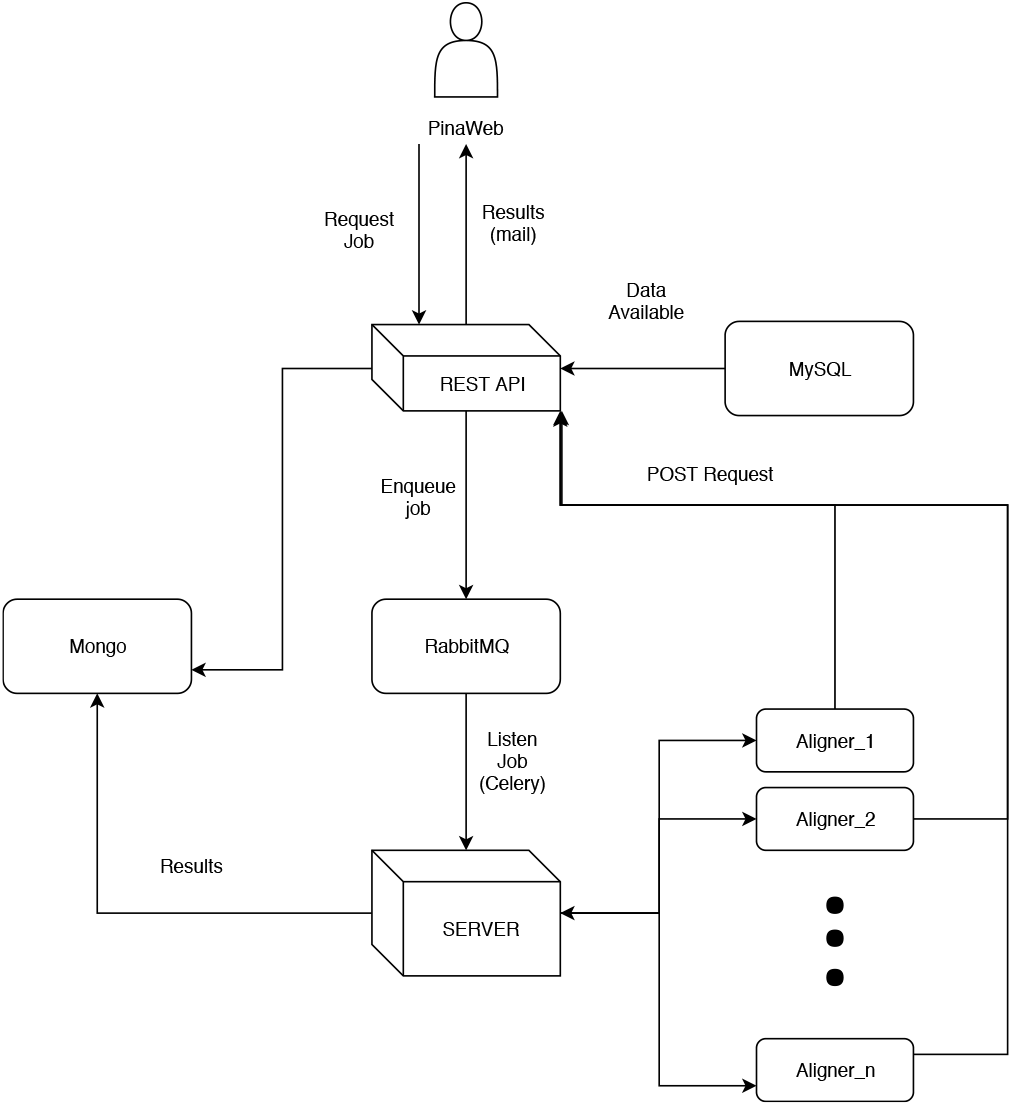
Diagram explaining the architecture of PINAWeb.

1. The user selects the alignment’s options: database, aligner, and networks.
2. The front-end sends a POST request to the REST API to create a job.
3. The REST API enqueues the job to the server using Celery and RabbitMQ as a broker.
4. The server computes the alignment and stores the results in a Mongo database.
5. The server calls back to the REST API with a POST request referencing the inserted result.
6. The REST API sends an email to the user with the results dashboard link.
7. The user can see the results via the results dashboard.

### Front-end web server

The user interacts with PINAWeb through the front-end. This front-end has a layer of css that uses *Bootstrap* to improve the user experience (UX) and user interface (UI), and it uses JavaScript to interact with the REST API through HTTP requests. Its main flow is then:

1. The front-end sends a GET request to the REST API to know which databases and aligners are available and it displays them at the corresponding interface selector.
2. When the user chooses a database, the front-end asks to the REST API which PPIN are available using another GET request and it displays them at the corresponding interface selector.
3. Finally, once the user has chosen the database, the aligner(s), and the networks, the front-end sends a POST request to the REST API with all this information and the user’s email and it shows a message to the user explaining that an email will be sent with the results.

### Dashboard

The outcome from the alignment can be visualized in a dashboard in order to help the user to compare the different alignments. It is implemented in Python 3.8 using the dash framework and deployed using Docker. The main flow is:

1. Get the *job id* from the path.
2. Request the results data to the REST API.
3. Computes the consensus for each protein.
4. Creates the html code to show the consensus and the algorithms results.
5. Returns the html.

### REST API

The main function of the REST API is to allow the user to create a job that the server will compute, and it is also in charge of sending the results to the user. It is implemented in Python 3.9 with all the libraries needed to create an asynchronous API (aiohttp, gunicorn, UVloop, aiomysql). The REST API has the following routes:

- */database* A GET request to ask which databases are available.
- */networks/ {database }*A GET request to ask which networks are available in the chosen database.
- */aligner* A GET request to ask which aligners are available.
- */create-job* A POST request to create a job in Celery which will be executed in the server.
- */finished-job* A POST request to create a task which will send the results of the job to the user.

The REST API uses REDIS to avoid computing the same job several times. So, if a user wants to compute an alignment which has been previously computed, the REST API simply sends the results to the user. Also if a job has been previously created but still not finished, the REST API adds the email of the user to the mailing list without creating a new job, so that once the alignment is computed, all users who have requested this alignment will receive the results.

### SERVER

The server is in charge of the computation of the alignments. It is implemented using essentially the same technologies as the REST API and works as follows:

1. A Celery worker is listening to a RabbitMQ queue waiting for a new job.
2. When a new job is created, the server gets all the parameters needed to compute the alignment from the job.
3. The server gets all the data needed to compute the alignment.
4. The server creates a system task to compute the alignment. In this way, is not needed to implement all aligners since we can use the binaries available. Given that all the architecture is implemented under Docker, if in the future we need any special system requirements to execute an aligner, we can easily add them to our system.
5. Once the alignment is computed, this is stored in the Mongo database and a POST request is sent to the REST API.

## Results and discussion

In this section we present our tool PINAWeb and present the results of several tests to discuss the tool’s usability. In order to explain how to use PINAWeb as well as the tool’s functionalities, we display here a query and its output as a running example. The query is to obtain the alignment for every aligner (AligNet, HubAlign, LGraal, Pinalog and Spinal) of two PPINs. As the input network, we considered a set of proteins from the Saccaromises scerevisiae and created a network. As the output network, we selected the Homo sapiens network from the STRING database, and we considered as edges the *Experimental score* with a threshold value of 800. PINAWeb computes the alignment of the input network to the output network for every selected aligner. When the computations are finished the user receives an email with a link to retrieve and analyze the obtained results.

### PINAWeb query webpage

Figure 2 shows PINAWeb’s query webpage whith the running example introduced as a query. The query webpage consists on different boxes where the user selects the aligners and introduce the input and the output networks as well as the email address where the results are sent. The information and usability of every box are explained below.

**Figure 2:**
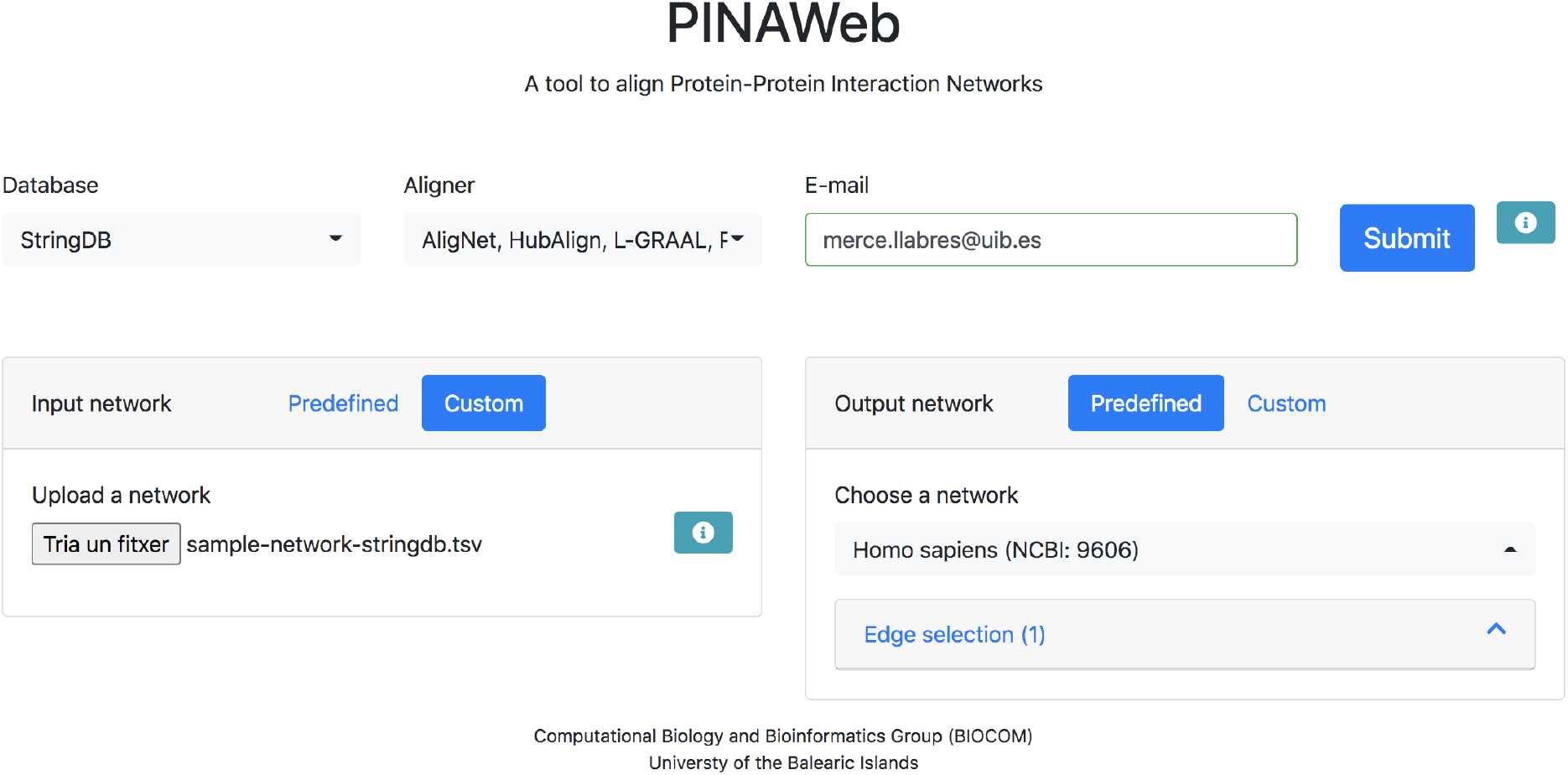
PINAWeb’s query example.

#### Database

PINAWeb retrieves from the STRING database the networks information as well as all the information needed to compute the requested alignments. Namely, the sequence similarity of every pair of proteins and the GO terms of every annotated protein. Therefore, the STRING database must be selected in this box.

#### Aligner

The available aligners are AligNet, HubAlign, L-GRAAL, PINALOG and SPINAL. The user can select all, only one, or several of them. The selected aligners are then displayed in the toolbox.

#### E-mail

The user has to provide an E-mail address where the tool sends the links to the results of every query.

#### Input network

In this box the input network of the query must be introduced or selected. In our running example it is a personalized network. To enter a personalized network click the button *Custom*. Then, it appears the option to upload a network from the user’s computer. In the information button on the right within this box it is explained the accepted format of the network to upload, as well as two examples of custom networks.

#### Output network

In this box the output network of the query must be introduced or selected. In our running example it is the *Homo sapiens* PPIN from the STRING database. To enter the network, the user must select the button *Predefined* and then look for the species name. To facilitate this search, the user can write the first letters of the species name to reduce the searching dataset. Next, in the *Edge selection* button, the user can choose all the edge types (score types) from the STRING database. For every selected edge type, a threshold must be introduced, and only the edges type whose weight is above the selected threshold in the STRING database will be included in the network.

### PINAWeb output-results

Once the requested alignments have been completed, PINAWeb sends an email to the address introduced in the query webpage with the job identifier. In this email the user will find a link to a webpage with the results. On top of this PINAWeb Results webpage the information is split in two menus. In the *Alignments info* menu all the single information of the requested alignments is provided (see Figure 3). Namely, the information is classified in the following sections:

**Figure 3:**
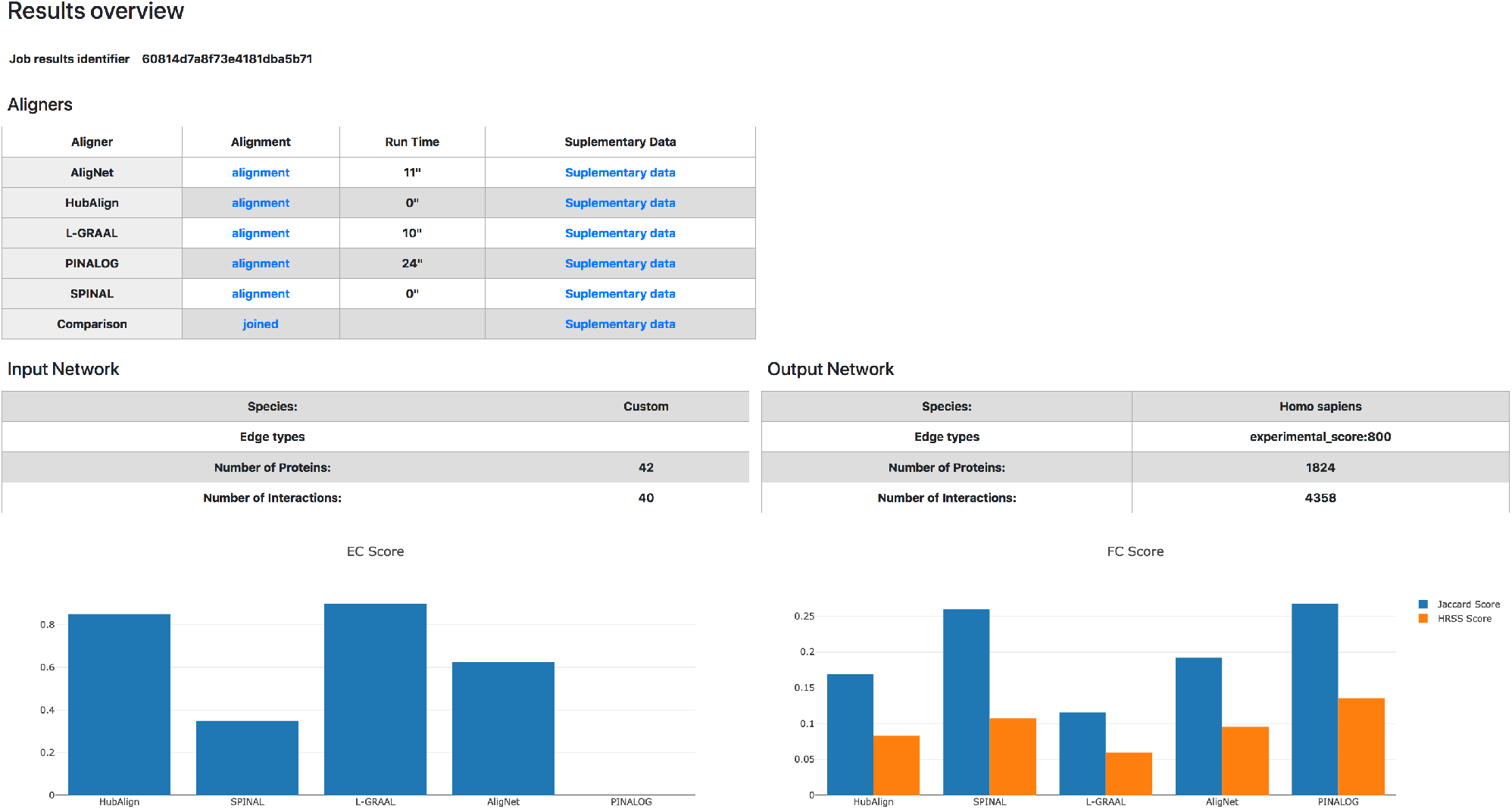
This figure displays the alignments information provided by PINAWeb. The quality measures of every alignment are presented in a barplot.

- **Aligners**: This table shows the aligner(s) for each alignment. In the the second column there is a link to the alignment results in a tsv file format. In the third column the run time in seconds is displayed for each aligner. In the last column, there is a link to a json file with all supplementary data of every alignment process. The json file also contains the unaligned nodes and edges, the non preserved nodes and edges and the annotated proteins. In the last row of the table, when more than one aligner was selected, PINAWeb provides a tsv file that merges the result of all requested alignments. Also, the corresponding json with all supplementary information is provided.
- **Input Network/Output Network**: The input and output networks requested by the user are shown in these two tables as well as their numbers of nodes and edges.
- **Alignment(s) quality measures**: PINAWeb summarizes the quality measures through a barplot diagram. The measures are: the *edge correctness ratio* (*EC*), that quantifies the amount of structure preserved by every alignment, and two functional coherence values, the *Jaccard Functional coherence value* and the *HRSS/BMA Functional coherence value* which assess the functional similarity of the aligned proteins by comparing their *Gene Ontology annotation*. More formally, let *G* = (*V, E*) and *G′* = (*V′, E′*) be two PPIN such that |*V* |*≤* |*V*| *′* and let *µ*: *V*→*V*′ be a mapping defining an alignment. The *edge correctness ratio* of *µ* is

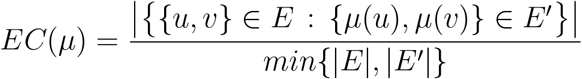

the *Jaccard Functional coherence value* of *µ* is

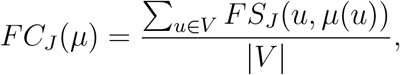

where the similarity score *FS*_*J*_ is defined by

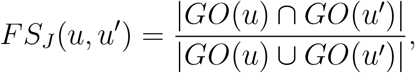

with *GO*(*u*) and *GO*(*u′*) the sets of GO annotations of the proteins *u* and *u′*, respectively, and the *HRSS/BMA Functional coherence value* of *µ* is

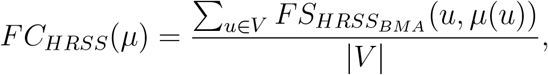

where the similarity score *FS*_*HRSS*_*BMA* is defined by

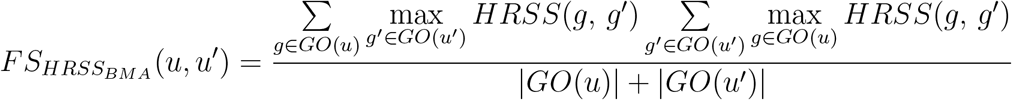

with *GO*(*u*) and *GO*(*u′*) the sets of GO annotations of the proteins *u* and *u′* respectively, and *HRSS*(*g, g′*) the *Hybrid Relative Specificity Similarity* of individual GO annotations. See (Clark & Kalita, 2014) and (Wu, Pang, Lin, & Pei, 2013) for a full description and discussion of the alignment quality measures introduced so far.

When more than one aligner is selected, in the *comparison* menu, PINAWeb provides a heatmap to visualize the comparison of the alignments’ results (see Figure 4). More precisely, every selected aligner is displayed on the left in the heatmap and all proteins from the input network are listed as columns. The user can select the number of proteins (columns) to visualize in the heatmap. The color corresponding to unaligned proteins is gray. The agreement level between the aligned proteins is represented through a color palette that ranges from light green to dark green. Dark green indicates a total agreement, meaning that all aligners have matched the protein to the same protein in the output network. Light green color means that all aligners matched the corresponding protein to a different one in the output network. Thus, a vertical column in dark green shows a consensus among all aligners. In addition, when pointing the cursor in the columns on the heatmap a window with the following information is shown: *Origin:* The protein name in the input network, *Target:* The protein name in the output network where the origin protein has been assigned by the aligner also displayed in the window.

**Figure 4:**
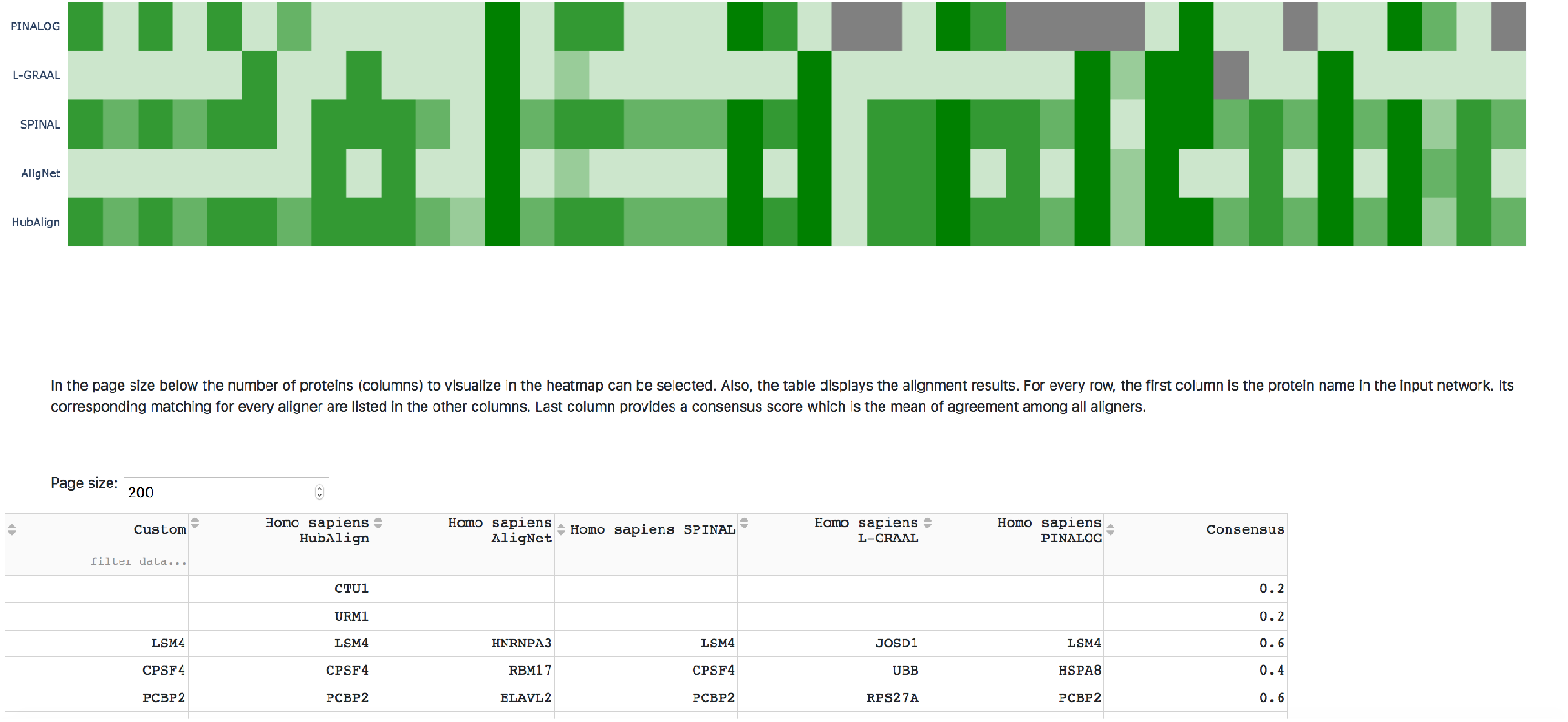
This figure shows the heatmap provided to compare the aligners results. The table displayed below the heatmap shows the matchings information.

### Experiments

We report here the experiments designed in order to show the tools usability. As a first experiment, we considered the PPINs of the organisms used in (Clark & Kalita, 2014) which consists of the PPIN of *M. musculus* (mus), *C. elegans* (cel), *D. melanogaster* (dme), *S. cerevisiae* (sce), and *H. sapiens* (hsa). The PPINs were selected from the STRING database with an experimental score of 800 as edge selection. In Table 1 we provide the networks information regarding their number of nodes and edges.

**Table 1:** Number of nodes and edges of the PPINs considered as input data in our first experiment.

|  | Nodes | Edges |
| --- | --- | --- |
| *C. elegans* | 509 | 1,149 |
| *M. musculus* | 751 | 1,443 |
| *D. melanogaster* | 1,224 | 1,933 |
| *H. sapiens* | 1,824 | 4,358 |
| *S. cerevisiae* | 2,775 | 13,269 |

We requested the alignments for every pair of PPINs and each aligner. Thus, we ended up with 50 alignments. As a result, PINAWeb performed all alignments and provided all the alignments’ information. We show the running time of every computation and each aligner in Table 2 below.

**Table 2:** Time in seconds of every alignment computation.

|  | AligNet | HubAlign | L-GRAAL | PINALOG | SPINAL |
| --- | --- | --- | --- | --- | --- |
| cel-mus | 32 | 0 | 117 | 3 | 11 |
| cel-dme | 30 | 0 | 314 | 9 | 10 |
| cel-hsa | 77 | 1 | 666 | 28 | 19 |
| cel-sce | 61 | 1 | 1193 | 401 | 75 |
| mus-dme | 52 | 1 | 553 | 10 | 4 |
| mus-hsa | 156 | 2 | 977 | 26 | 13 |
| mus-sce | 90 | 2 | 2444 | 447 | 63 |
| dme-hsa | 127 | 7 | 2148 | 37 | 20 |
| dme-sce | 104 | 6 | 3607 | 451 | 35 |
| hsa-sce | 203 | 28 | 3604 | 530 | 113 |

**Table 3:** Link to the results obtained in the tests.

|  |  |
| --- | --- |
| cel-mus | https://biocom.uib.es/util-aligner-results/607944acf9ef164d857915f7 |
| cel-dme | https://biocom.uib.es/util-aligner-results/6075b725f9ef164d85791504 |
| cel-hsa | https://biocom.uib.es/util-aligner-results/6075b620f9ef164d857914d6 |
| cel-sce | https://biocom.uib.es/util-aligner-results/6075bb65f9ef164d8579154b |
| mus-dme | https://biocom.uib.es/util-aligner-results/6075b767f9ef164d8579151b |
| mus-hsa | https://biocom.uib.es/util-aligner-results/6075b6f3f9ef164d857914ed |
| mus-sce | https://biocom.uib.es/util-aligner-results/6075be1ff9ef164d85791579 |
| dme-hsa | https://biocom.uib.es/util-aligner-results/6075bc5bf9ef164d85791562 |
| dme-sce | https://biocom.uib.es/util-aligner-results/6075b81ff9ef164d85791534 |
| hsa-sce | https://biocom.uib.es/util-aligner-results/607d48768f73e4181dba5a8f |
| Test 2 | https://biocom.uib.es/util-aligner-results/6082d6628f73e4181dba5b9f |

As we can observe there, HubAlign is the fastest aligner followed by SPINAL while L-GRAAL is the slowest. AligNet performed each computation in less than 4 minutes and PINALOG was faster than AligNet in most computations except those with S. cerevisiae, probably due to its large number of edges. Nevertheless, in one hour PINAWeb reported all the requested alignments, which reinforces the correct election of the aligners.

Regarding the alignments themselves, the highest quality measures were obtained in the alignment of M. musculus and H. sapiens. This may reveal that, as expected, when the networks to be aligned are similar (closer species), the aligners perform better. Indeed, Figure 5 shows the hierarchical clustering considering the euclidean distance and complete linkage of the consensus alignment scores. We can observe that the cluster of M. musculus and H. sapiens is clearly separated from the other species.

**Figure 5:**
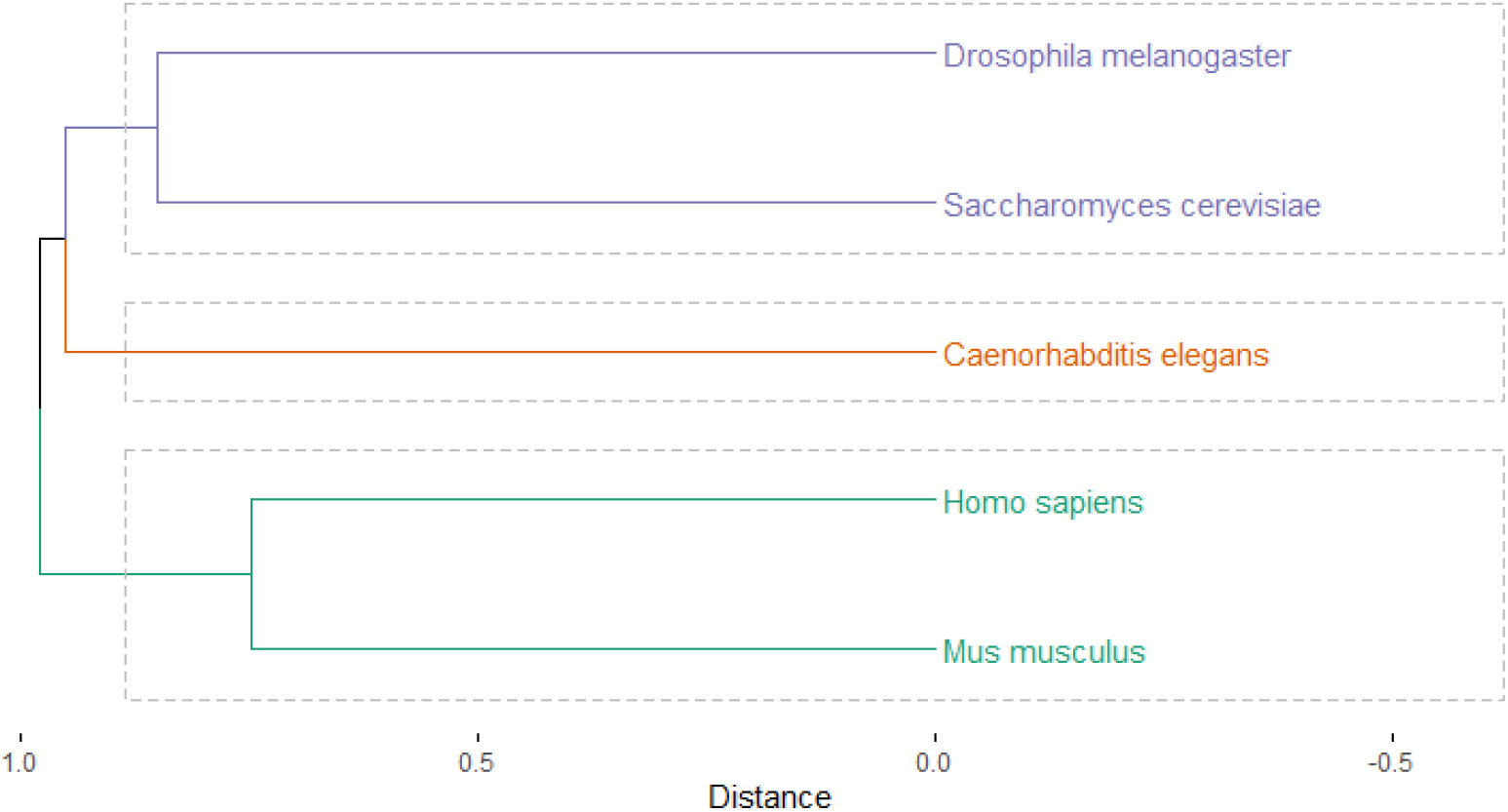
Hierarchical clustering of the consensus alignment score in the first experiment.

As a second experiment, in order to evaluate the consensus information provided by PINAWeb, we designed the following test: we considered the same PPIN of H. sapiens as in the previous test and we randomly deleted some nodes and edges. Thus, we synthetically created two PPINs as subnetworks of the H. sapiens PPIN such that they have 58 and 67 proteins respectively, and they share 45 of them. The networks have 64 and 76 edges respectively. Then, we requested the alignment of this pair of subnetworks for every alginer. As a result, we obtained that, except PINALOG, all aligners properly matched the shared proteins. That is, all aligners mapped every protein present in both networks to itself. The consensus alignment in this case was 0.8. Also the edge correctness ratio and the Jaccard functional coherence value were above 0.8. Therefore we conclude that the aligners obtained a considerable agreement in this test.

In the table below we provide the links to all the alignments and comparison results obtained in the experiments reported here.

## Conclussion

In this paper we introduce PINAWeb, a user-friendly web-based tool aimed to obtain the alignments produced by the aligners AligNet, HubAlign, L-GRAAL, PINALOG and SPINAL, thus avoiding the tiresome work of data processing, installation and computations for every aligner. Alignments are performed either on networks retrieved from the STRING database or from the user’s own data. In addition, PINAWeb yields a heatmap visualization to compare the aligners performance and provides a table with the matching results as well as a consensus score, when more than one aligner is selected. Additionally, the output also provides a json file with the networks information (nodes and edges) as well as the GO terms of the involved proteins, the unaligned nodes and edges, the non preserved nodes and edges and the annotated proteins.

Two tests has been carried on to show the tool’s usability. In the first test five networks were selected and the “all against all” network pairs alignment was performed for each aligner. Thus, we ended up with 50 alignments. PINAWeb produced all the requested alignments in an average time of 5 minutes. In addition, the comparison of the results revealed that all aligners reach a higher agreement on similar networks. To reinforce this conclusion, in the second test a pair of subnetworks from H. Sapiens was aligned resulting, indeed, in a very high agreement among the aligners. Therefore, we encourage the researchers to use PINAWeb to improve their performance when analyzing their experiments results.

## Acknowledgements

We acknowdledge the Ministerio de Ciencia e Innovación (MCI), the Agencia Estatal de Investigación (AEI) and the European Regional Development Funds (ERDF) for its support to the project PGC2018-096956-B-C43.

